# Polarity sensitive probes for super resolution STED microscopy

**DOI:** 10.1101/107334

**Authors:** E Sezgin, F Schneider, V Zilles, E Garcia, D Waithe, A S Klymchenko, C Eggeling

## Abstract

The lateral organization of molecules in the cellular plasma membrane plays an important role in cellular signaling. A critical parameter for membrane molecular organization is how the membrane lipids are packed (or ordered). Polarity sensitive dyes are powerful tools to characterize such lipid membrane order, employing for example confocal and two-photon microscopy. The investigation of potential lipid nanodomains, however, requires the use of super resolution microscopy. Here, we test the performance of the polarity sensitive membrane dyes Di-4-ANEPPDHQ, Di-4-AN(F)EPPTEA and NR12S in super resolution STED microscopy. Measurements on cell-derived membrane vesicles, in the plasma membrane of live cells, and on single virus particles show the high potential of these dyes for probing nanoscale membrane heterogeneity.

## Introduction

The lateral organization of molecules in the cellular plasma membrane has strong influence on cellular functions. Lipid-lipid and lipid-protein interactions facilitate the segregation of plasma membrane molecules, for example into clusters or nano-domains which constitute catalytic platforms for a myriad of activities such as cellular signaling (1). Lateral heterogeneity in lipid order, i.e. how the packing of lipids changes over space, and how this is involved in molecular segregation, is particularly of interest since membrane order may modulate protein functionality (2-5). Saturated lipids can be ordered relatively more tightly in contrast to unsaturated lipids that yield less ordered (i.e. more disordered) membranes. Polarity sensitive fluorescent probes such as Laurdan are useful tools to study such lipid order (6-8). These dyes change their emission spectrum depending on the environmental conditions; for example, they exhibit a relatively more red-shifted fluorescence spectrum in more polar solvents (9, 10). More tightly packed ordered lipid environments accommodate less amount of water molecules and are therefore relatively more nonpolar than disordered environments (11). In turn, the fluorescence emission spectrum of the polarity sensitive dyes is more blue-shifted in ordered membrane environments, which can be used to quantify the molecular ordering and to visualize lateral heterogeneity in membrane order of cellular membranes (12). These probes have been used in combination with confocal or multi-photon microscopy (13-15); however, the diffraction-limited spatial resolution of these techniques does not allow observation and full characterization of nanodomains/clusters in the plasma membrane. A remedy to this barrier is super-resolution optical microscopy such as STED microscopy (16, 17). Although STED microscopy has recently been employed to study nano-scale plasma membrane organization (18-22), the combination of this technique with polarity sensitive probes has not yet been realized. The institution of traditional polarity sensitive probes such as Laurdan and Prodan suffers from their low photostability and near-UV excitation, already in conventional microscopy (13). Recently developed probes such as Di-4-ANEPPDHQ (23), Di-4-AN(F)EPPTEA (24) and NR12S (25) (Figure 1) show higher brightness and red-shifted operating range, thus these probes can be suitable for STED imaging. Here, we test the performance of these polarity sensitive membrane probes in STED microscopy. Using cell-derived membrane vesicles, live CHO cells and virus particles, we highlight their potential for studying lateral differences in plasma membrane order with sub-diffraction spatial resolution.

**Figure 1.**
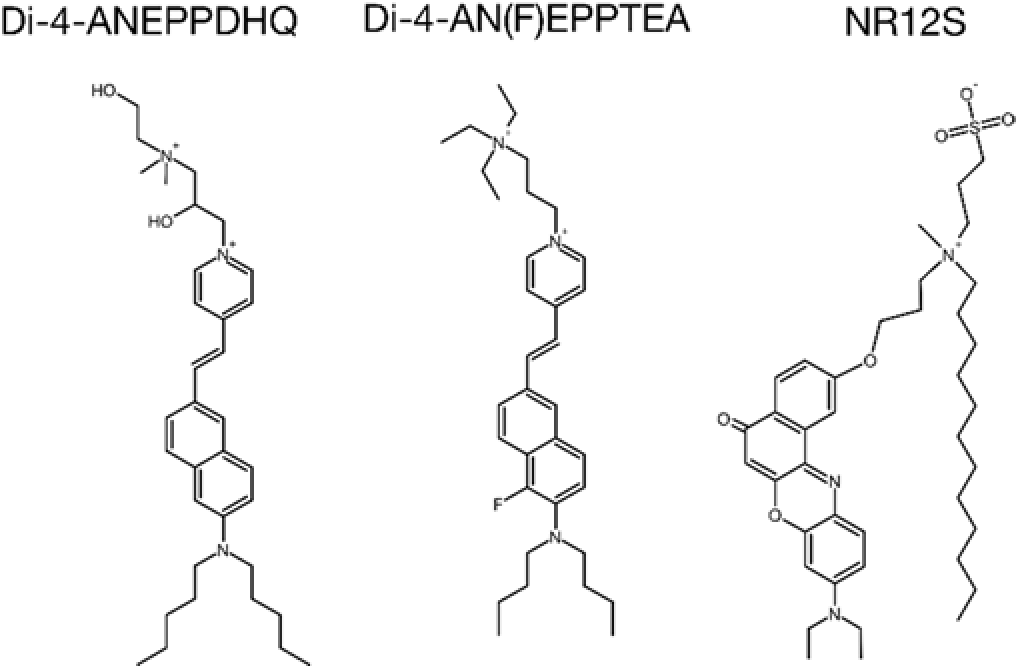
Structures of the polarity sensitive membrane dyes employed in this study; Di-4-ANEPPDHQ, Di-4-AN(F)EPPTEA and NR12S.

## Materials and Methods

### Materials

Laurdan and Di-4-ANEPPDHQ were purchased from ThermoFisher. Di-4-AN(F)EPPTEA (24) was a kind gift from Prof. Leslie Loew. F2N12SM, FC12SM, PA, and NR12S were made as previously described (25-28). Abberior Star Red-labelled 1,2-Dipalmitoyl-sn-glycero-3-phosphoethanolamine (DPPE) was obtained from Abberior (Gottingen, Germany).

### Cells and GPMV preparation

CHO cells were maintained in F12 DMEM (Sigma) medium supplemented with 10% FBS (Sigma) and 1% L-glutamine (Sigma). Cells were seeded on 35 mm petri dishes (plastic dishes for GPMV experiments (Nunc), and glass-bottom dishes (Ibidi) for cellular imaging experiments) three days before the experiments. For GFP transfection, the cells were transfected with cytoplasmic GFP plasmid in the second day using Turbofect (Thermofisher) transfection agent. GPMVs were prepared using 25 mM paraformaldehyde (PFA, Sigma) and 2 mM or 10 mM Dithiothreitol (DTT, Sigma) in GPMV buffer for non-phase-separated and phase-separated vesicles, respectively, as detailed in (29). GPMVs were immobilized as shown in Supplementary Figure S1 for imaging as detailed in ref (30). Cells were labelled with the polarity sensitive membrane probes at 1 µg/ml final probe concentration in phosphate buffered saline (PBS) for 5 minutes before imaging. Live cell imaging was performed in L-15 medium (Sigma) without phenol red. GPMVs were labelled with 0.1 µg/ml final probe concentration after before imaging. All imaging experiments were performed at room temperature.

### Virus particle preparation

Virus particles were prepared as detailed in ref (31). In brief, they were generated from the tissue culture supernatant of 293T cells co-transfected using polyethyleneimine (PEI) with 14 μg pCHIV per 10 cm dish. Tissue culture supernatants were harvested 48 h after transfection, cleared by filtration through a 0.45 μm nitrocellulose filter, and particles were purified by ultracentrifugation through 20% (w/w) sucrose cushion at 70,000 g (avg.) for 2 hr at 4 °C. Virus containing gradient fractions were diluted in PBS and pelleted at 70,000 g (avg.) for 2 hr at 4 °C. Particles were resuspended in ice-cold 20 mM HEPES/PBS pH 7.4, snap frozen and stored in aliquots at −80 °C. All ultracentrifugation steps were performed in SW 41 Ti rotor. For microscope measurements, purified virus particles were adhered to poly-L-lysine (Sigma) coated glass cover slips for 30 min. Cover slips were blocked using 2% bovine serum albumin (BSA) (Sigma)/PBS for 15 min. Particles were stained for polarity sensitive dyes with 10 µg/ml final concentration by 30 min incubation. After staining, the particles were washed and mounted in PBS, followed by microscopy analysis. All steps were carried out at room temperature.

### Spectral Imaging

Spectral imaging was performed on a Zeiss LSM 780 confocal microscope equipped with a 32-channel GaAsP detector array. Laser light at 405 nm was selected for fluorescence excitation of Laurdan and 488 nm for the other polarity sensitive probes. The lambda detection range was set between 415 and 691 nm for Laurdan, and between 495 and 691 nm for the other dyes. The images were saved in .lsm file format and then analyzed by using a custom plug-in compatible with Fiji/ImageJ, as described in (32).

### Confocal and STED Imaging

All images were acquired using the Leica SP8 STED microscope (Leica Microsystems, Mannheim, Germany) equipped with a HC PL APO C S2 100x / 1.40 oil objective (Leica Microsystems), a pulsed (80 MHz) white light laser (WLL), and a continuous-wave (CW) 592 nm, CW 660 nm, and pulsed 775 nm STED lasers. We selected 488 nm for excitation and 775 nm for STED. The master laser power for the WLL was 70 % and for the STED laser kept at 75 %. The filter used was Notch Filter (NF) 488/561/633. Before each measurement the beam alignment of the WLL and the STED laser was checked. The alignment was performed with the 592 nm STED laser (reference laser). The laser power of the WLL laser was in the range of 10–25 %, resulting in approximately 10-20 µW at the sample, while the maximum employed STED laser power was 75 % (≈100 mW) at the sample. The emission was detected using hybrid detectors. When two wavelengths were needed, 530-570 nm interval and 650-690 nm intervals were selected. Gating between 1.5-6 ns was applied. Two image sequences were taken; the first sequence without the STED laser on and the second sequence with 75% STED-laser power. The first sequence provided the standard confocal image whereas the second sequence the high resolution STED microscopy image. The scanning speed was set to 1800 Hz line frequency and a line-averaging over 4 lines was applied. All images had a zoom factor of 10 and the same image size. The pixel size was optimized for STED image and kept the same for confocal.

### Calculation of membrane width from microscopy images

The recorded images were analyzed with the macro program “one line” in the software FiJi to determine the full-width-at-half-maximum (FWHM) values of membrane width in the microscope images. The FWHM analysis was carried out for multiple intensity line profiles across the imaged membrane. A bulk FWHM measuring algorithm was designed and written using ImageJ macro language (https://github.com/dwaithe/generalMacros/tree/master/FWHM%20bulk%20measure). Using the ImageJ interface the user defines a line using the “Segmented line” tool which follows the contour of the vesicle to be measured. The algorithm is then run and interpolates along the user-defined line and calculates points at regular intervals (3 pixel gap). At each interpolated point on the line perpendicular lines are drawn 40 pixels in length and centered on the interpolated line. Along each of these perpendicular lines the intensity values are sampled and a Gaussian curve is fit using the ImageJ curve fitting plugin (see Supplementary Figure S2). The parameters of each curve are then output and the FWHM calculated for each curve (2x√(2xlog(2)*σ), where σ is the standard deviation of the Gaussian fit). The extracted FWHM-values were averaged over at least 10 images. Confocal and STED images were treated in the same way.

### Results and Discussion

In our experience, a polarity sensitive membrane probe adequate for advanced imaging (such as time-lapse or super resolution imaging) of lateral plasma membrane heterogeneity should have (i) a high sensitivity to membrane lipid packing, (ii) a high photostability, and (iii) an appropriate narrow emission spectrum to allow its combination with other fluorescent probes for labeling membrane proteins. Such properties have been indicated for recently developed probes such as Di-4-ANEPPDHQ, Di-4-AN(F)EPPTEA and NR12S (13, 23-25, 27, 28) (Figure 1).

We first compared the performance of the probes with respect to their sensitivity to differences in lipid order. For this purpose, we employed phase-separated model membranes that feature coexistence of a disordered and an ordered membrane phases. To mimic conditions in the plasma membrane as close as possible, we used cell-derived giant plasma membrane vesicles (GPMVs) (29, 33, 34). The differences in lipid packing between disordered and ordered phases has been shown to be much lower and more realistic in GPMVs than in purely artificial phase-separated model membrane systems such as giant unilamellar vesicles (GUVs) (2, 35). Therefore, GPMVs are more accurate tools to test the performance of the probes when their applicability in cellular context is evaluated. Figure 2A shows confocal images for different spectral ranges of the equatorial plane of phase separated GPMVs derived from the plasma membrane of CHO cells and labeled with the probe NR12S using spectral imaging (32). These images demonstrate differences in the emission spectra of the probes in ordered and disordered phases. The extent of the spectral shift is an adequate indicator on how sensitive the probes report on differences in plasma membrane order. For quantification of the observed spectral shift, we studied the fluorescence emission spectra of the investigated polarity sensitive probes in the ordered and disordered phases of GPMVs, with Laurdan as a control (Figure 2B-E). While still lower than of Laurdan (~50 nm, Figure 2B) (9), all probes showed a clearly visible spectral shift of >30 nm. In contrast, other recently developed polarity sensitive probes F2N12SM (27), FC12SM (27) and PA (28) showed much lower spectral shift <15 nm (Supplementary Figure S3). We concluded that Di-4-ANEPPDHQ, Di-4-AN(F)EPPTEA and NR12S are sensitive enough to distinguish ordered and disordered environments in cellular membranes.

**Figure 2.**
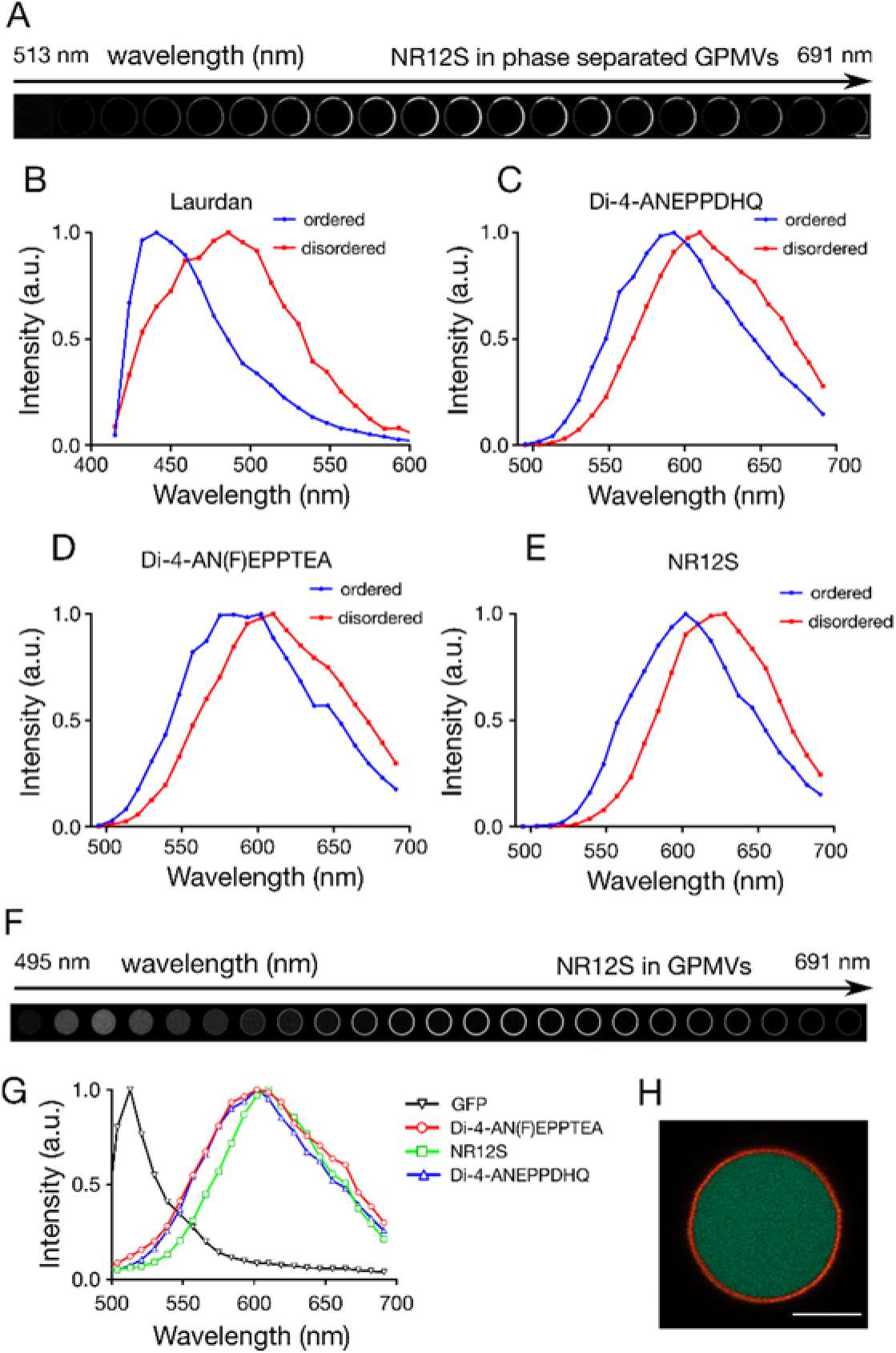
Spectral imaging of the polarity sensitive probes in CHO cell-derived GPMVs. A) Representative confocal images of the equatorial plane of a phase-separated GPMV stained with NR12S for subsequent 8 nm wide spectral windows between 495 and 691 nm, indicating the emergence of fluorescence maxima in different parts. B-E) Exemplary fluorescence emission spectra of B) Laurdan, C) Di-4-ANEPPDHQ, D) Di-4-AN(F)EPPTEA and E) NR12S in the disordered (red) and ordered (blue) phases of the phase separated GPMVs. F) Representative confocal images of the equatorial plane of a non-phase separated GPMV derived from CHO cells transfected with cytoplasmic GFP and membrane-stained with NR12S for subsequent 8 nm wide spectral windows between 495 and 691 nm, indicating a spectral separation of fluorescence signal from the GPMV body (GFP) and membrane (NR12S). G) Exemplary fluorescence emission spectra of GFP (black) and the polarity sensitive dyes in non-phase separated GPMVs (Di-4-ANEPPDHQ (blue), Di-4-AN(F)EPPTEA (red) and NR12S (green)). H) Fluorescence signal from panel F for the spectral ranges between 500 and 530 nm (green) and 570 to 691 nm (red), indicating clear separation of the GFP and NR12S signal, respectively. Scale bars 5 µm.

Protein functionality may be modulated by the state of the lipid order of its immediate membrane environment. Therefore, it is crucial to be able to determine lipid packing around specific proteins, which requires simultaneous labelling and observation of the proteins (e.g. using green fluorescent protein, GFP) and of polarity sensitive dyes. For distinguishing both, their spectra have to be well separated, i.e. the spectra of the polarity sensitive dyes have to be narrow enough not to significantly overlap with the fluorescence emission spectrum of GFP. We therefore performed spectral imaging on non-phase separated GPMVs derived from CHO cells transfected with cytoplasmic GFP (i.e. the GPMVs were filled with GFP) and membrane-labelled with the selected polarity sensitive dyes. Here, we chose non-phase separated GPMVs, since we now aimed at separating two homogeneous features (membranes and body). The membrane lipid packing of non-phase separated GPMVs is in-between that of ordered and disordered environments in phase-separated GPMVs (32). Figure 2F shows confocal images for different spectral ranges of the equatorial plane of such a GPMV, indicating a clear spectral separation of fluorescence signal from the GPMV body (GFP) and membrane (NR12S). The possibility of separating fluorescence emission from GFP and the polarity sensitive probes is indicated in Figure 2G, which depicts the fluorescence emission spectra reconstructed from the spectral images of the GPMV bodies (GFP) and membranes (polarity sensitive probes). In all cases (Di-4-ANEPPDHQ, Di-4-AN(F)EPPTEA and NR12S), a significant separation between both spectra is observed. With appropriate filters (such as a 500-530 nm bandpass and a 570 nm longpass filters for GFP and the polarity sensitive dyes, respectively), the fluorescence signals from GFP and the polarity sensitive dyes can be clearly separated as highlighted in Figure 2H, which plots the fluorescence signal from Figure 2F for the spectral ranges between 500 and 530 nm (green) and 570 to 691 nm (red).

Next we tested the performance of the polarity sensitive probes in STED microscopy. We again incorporated the probes into CHO cell-derived GPMVs and imaged the equatorial planes with confocal and STED microscopy using 488 nm and 775 nm laser light for excitation and depletion, respectively. Again we chose non-phase separated GPMVs, since we aimed here at imaging the width of the membrane, which we aimed to have homogeneous. A representative confocal and STED image of the same GPMV stained with Di-4-AN(F)EPPTEA is shown in Figure 3A. Obviously, the membrane appears thicker in the confocal compared to the STED microscopy image, as highlighted by intensity line profiles across the membrane (Figure 3B). Since the thickness of the cell membrane (≈8 nm) is well below the expected spatial resolution of both microscopy modes, the thickness of the imaged membrane is a direct indicator of the achieved spatial resolution. In order to give an estimate of the increase in spatial resolution between confocal and STED microscopy recordings, we determined the width (full width at half maximum (FWHM)) of the respective membrane images using a custom FiJi macro (see Methods and Supplementary Figure S2), and divided the FWHM values obtained for the confocal images by those obtained for the STED microscopy images; values >1 indicate an increase in resolution and the larger the values the better the improvement. We obtained similar ratios (≈3, 70-90 nm spatial resolution) for all of the tested polarity sensitive dyes, which were similar to the FWHM values obtained for confocal and STED microscopy images of GPMVs stained with a KK114-labeled phospholipid (Figure 3C). KK114 (or Abberior Star Red) is a well-established dye for STED microscopy (36) and the KK114-stained phospholipid has successfully been employed in previous STED microscopy studies (20, 22).

**Figure 3.**
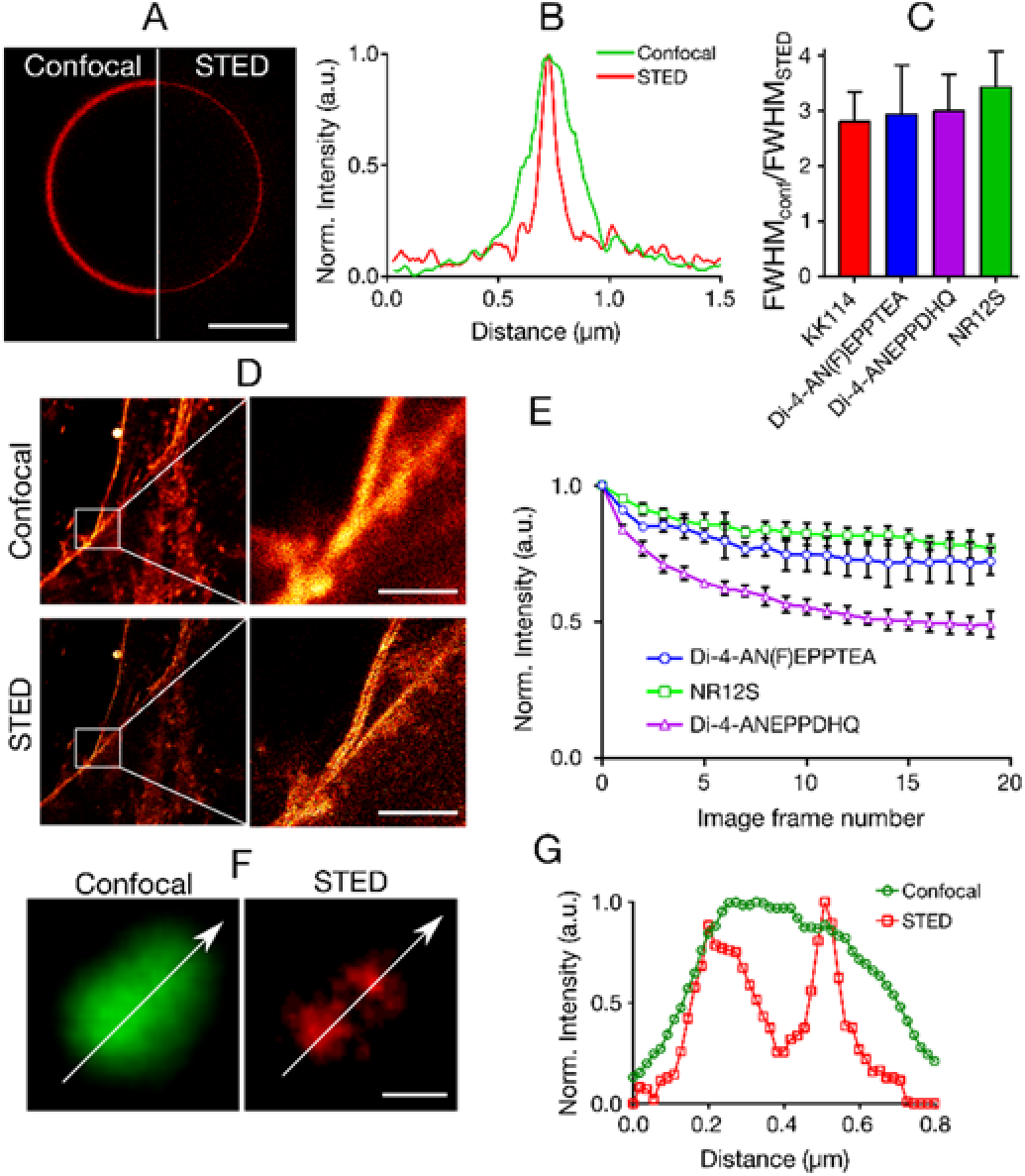
STED microscopy of polarity sensitive membrane probes. A) Representative confocal (left) and STED (right) microscopy images of the equatorial plane of the same CHO cell-derived GPMV stained with Di-4-AN(F)EPPTEA. Scale bar 10 µm. B) Intensity line profiles through the membrane as obtained from the confocal (green) and STED (red) images of panel A. C) Average ratio of FWHM values obtained from the line profiles through the membranes in confocal and STED microscopy images of the CHO cell-derived GPMVs stained with Di-4-ANEPPDHQ (purple), Di-4-AN(F)EPPTEA (blue), NR12S (green) and a KK114-labelled phospholipid (red). Error bars are standard deviation from at least 10 measurements. D) Representative confocal (upper) and STED (lower) microscopy images of the same CHO cells labelled with Di-4-AN(F)EPPTEA (left: overviews; right: zoom-ins into marked areas), highlighting the improvement in spatial resolution by separation of the plasma membranes of two adjacent cells. Scale bars 10 µm. E) Decrease in fluorescence signal for subsequent STED microscopy recordings of the same region of interest of CHO cells stained with Di4-ANEPPDHQ (purple), Di-4-AN(F)EPPTEA (blue), NR12S (red) (signal relative to first recording, average and error bars as standard deviation from at least 10 regions), indicating sufficient photostability of all probes. F) Representative confocal (green) and STED (red) microscopy images of two nearby HIV particles membrane-labelled with Di-4-AN(F)EPPTEA and G) intensity profiles along the lines marked in the images. Scale bar 100 nm.

To confirm the suitability of the polarity sensitive membrane probes for STED microscopy, we labelled the plasma membrane of live CHO cells with Di-4-AN(F)EPPTEA and imaged it using confocal and STED microscopy. Figure 3D shows representative side-by-side images in both modes, highlighting the clear improvement in spatial resolution when employing STED microscopy. In addition, time-lapse STED microscopy imaging of the same part of the cells incubated with the three fluorescent probes showed their reasonable photostability, with less than 50% (Di-4-ANEPPDHQ) and < 25% (Di-4-AN(F)EPPTEA and NR12S) loss in fluorescence signal after 20 subsequent images (Figure 3E).

The efficiency of a STED microscope depends on the fluorescence depletion efficiency by the STED laser light, which roughly scales with the position of the STED laser wavelength at the fluorescence emission spectrum, i.e. the further away the STED laser wavelength from the fluorescence emission maximum the lower the fluorescence depletion efficiency (16, 17). Therefore, one may expect a worse performance of the polarity sensitive dyes in STED microscopy when incorporated in more ordered membrane environments, where the emission spectrum is blue-shifted. To test this, we next explored the possible use of the polarity sensitive dyes for imaging and separating sub-diffraction sized HIV virus particles. The HIV membrane environment is characterized by a very high membrane order, much larger than in the cellular plasma membrane (37). We labeled HIV particles (≈100-150 nm in diameter) with Di-4-AN(F)EPPTEA and imaged them after immobilization to poly-L-lysine coated microscope cover glass. Figure 3F and 3G highlight the separation of nearby virus particle in STED microscopy imaging but not in the confocal mode, despite the more blue-shifted spectra. Clearly, the extend of the spectral blue shift (<30nm) is too low to significantly compromise the STED microscopy performance.

## Conclusion

We have here shown the compatibility of the polarity sensitive membrane dyes Di-4-ANEPPDHQ, Di-4-AN(F)EPPTEA and NR12S for super resolution STED microscopy. Using cell derived GPMVs, live CHO cells and HIV particles, we highlighted for all three probes (i) a sufficient sensitivity for distinguishing disordered and ordered environments in cellular membranes, (ii) the possibility to spectrally separate their fluorescence signal from GFP signal, (iii) the successful application in STED microscopy imaging of cellular membranes achieving a spatial resolution of around 70-90 nm in this case, and (iv) a sufficiently large photostability allowing for time-lapse imaging of living cells. These features provide a starting point for future measurements of lateral heterogeneity in lipid order in the plasma membrane of living cells with relation to the organization of e.g. GFP-labelled proteins. This may shed new light on the existence and function of possible lipid nanodomains of higher molecular order. While these domains are expected to be even smaller than the spatial resolution of 70-90 nm achieved so far, we expect with further optimization of the setup (such as laser intensity and wavelength) to improve the resolution further. Institution of a spectral detector for spectral STED imaging will then give the potential to directly observe membrane nanodomains of higher lipid order and solve their long-standing mystery. This study also sets a guideline for the development of future polarity sensitive probes suitable for advanced super-resolution imaging.

## Author Contributions

ES, FS and VZ carried out experiments. EG helped with microscopy recordings. DW wrote the analysis software and helped analysing the data. ES and CE conceived the study and wrote the manuscript with help from all authors. All authors contributed in discussing the data, experiments and the manuscript.

## Acknowledgements

We would like to thank Silvia Galiani for help on the STED microscope, Jakub Chojnacki and Pablo Carravilla for virus preparation, and Christoffer Lagerholm for help on the microscopes in the Wolfson Imaging Centre. Leslie Loew and Yosuke Niko are acknowledged for providing Di-4-AN(F)EPPTEA and PA probes, respectively. E.S. is supported by EMBO long term (ALTF 636-2013) and Marie Skłodowska-Curie Intra-European Fellowships (MEMBRANE DYNAMICS-627088). This work is supported by the Wolfson Foundation (ref. 18272), the Medical Research Council (MRC, grant number MC_UU_12010/unit programmes G0902418 and MC_UU_12025), MRC/BBSRC/ESPRC (grant number MR/K01577X/1), and the Wellcome Trust (grant ref. 104924/14/Z/14).

